# PARG inhibition induces nuclear aggregation of PARylated PARP1

**DOI:** 10.1101/2023.08.12.553088

**Authors:** Sateja Paradkar, Julia Purcell, Annie Cui, Sam Friedman, Ranjini Sundaram, Ryan Jensen, Ranjit Bindra

## Abstract

PARG inhibitors are currently under clinical development for the treatment of DNA repair-deficient cancers, however, their precise mechanism of action is still unclear. Here we report that PARG inhibition causes increased nuclear PARylated PARP1 that limits PARP1 chromatin binding in response to DNA damage. This PARylated PARP1 accumulates as aggregates at sites distinct from the site of DNA damage, leading to the mis-localization of PARP1. Additionally, these aggregates are formed through PAR chains as abrogating PARP1 catalytic activity prevents their formation. Finally, these PARP1 nuclear aggregates persist long-term and are associated with cleaved cytoplasmic PARP1, a cell death hallmark, which ultimately leads to a non-apoptotic form of cell death. Overall, our data uncovers a novel mechanism of PARG inhibitor cytotoxicity, which will inform ongoing clinical studies.

## Introduction

Poly(ADP-ribose) polymerase 1 (PARP1) is a central player in multiple biological processes including DNA repair^1^. In response to DNA damage, PARP1 binds to damaged DNA and synthesizes long chains of repeating ADP-ribose units using nicotinamide adenine dinucleotide (NAD^+^) as a substrate. PARP1 catalyzes the transfer of these poly(ADP-ribose) (PAR) chains onto target proteins, neighboring histone tails, and on PARP1 itself termed as autoPARylation^2-4^. PAR chains deposited at histones allow for chromatin relaxation which also facilitates the recruitment of DNA repair factors to the damage site. These PAR chains can contain as many as 200 ADP-ribose units^5^. PARP1 can also produce branched PAR chains, extending the size and structure of this modification^5^. Poly(ADP-ribose) glycohydrolase (PARG) is the primary enzyme responsible for the degradation of linear and branched PAR chains following DNA repair. PARG cleaves the ribose-ribose linkages in PAR chains into individual ADP-ribose units, allowing the efficient recycling of PAR. Owing to the critical role of PARP1 in DNA repair, multiple PARP inhibitors (PARPis) have been developed and approved to selectively target DNA repair-deficient cancers. These include cancers with mutations in key DNA repair proteins such as BRCA1/2, which occur in 50-70% of breast cancers and 5-15% of ovarian cancers^6,7^. While PARPis have provided therapeutic benefit in these cancers, clinical evidence demonstrates that resistance to these drugs emerges rapidly^8,9^.

To overcome the limitations of PARPis, researchers have been developing PARG inhibitors (PARGis) for the treatment of DNA repair-deficient cancers particularly in instances of PARPi resistance. Previous reports have suggested that PARPis and PARGis could target different subsets of ovarian cancer patient populations depending on underlying replication vulnerabilities^10^. Given this, there are currently multiple PARGis in various stages of pre-clinical and clinical development. As these drugs are developed, it becomes imperative to understand the precise mechanism of how these inhibitors disrupt DNA repair and related processes prior to their use in patients. PARGis have been shown to cause increased PARylation at DNA damage sites and consequently an increased retention of proteins recruited in response to PAR at these damage sites^11-13^. Additionally, PARG inhibitors have been shown to cause increased DNA damage, increased sensitivity to DNA damaging agents, and increased replication fork defects^11,12,14,15^. However, the precise mechanism underlying these phenotypes still remains to be determined. As PARG is a close cooperator of PARP1, we hypothesized that PARG inhibition could be disrupting these cellular processes through its effects on PARP1 function.

Here we report a novel mechanism of cytotoxicity as a result of PARG inhibition. Using laser microirradiation and live cell imaging, we demonstrate that PARG inhibition leads to the accumulation of nuclear PARylated PARP1 aggregates at distinct sites from the original site of damage. We observe that these aggregates persist long after PARP1 dissociation from DNA in untreated conditions. By testing PARP1 catalytic variants in the same live cell imaging experiments, we show that PARP1 activity is necessary for the formation of these aggregates. Finally, we show that these aggregates are associated with increased cell death in a BRCA-proficient and deficient cell line. Overall, this finding provides the framework for understanding the effects of PARG inhibition on PARP1-mediated nuclear functions such as DNA repair, replication, and cell death.

## Results

### PARG inhibition reduces PARP1 chromatin binding in response to DNA damage

Owing to the central role of PARG in PARP1-mediated DNA repair, we first began by studying the effect of PARG inhibition on PARP1 chromatin recruitment in response to DNA damage. Previous reports have observed changes in PARP1 levels in the nuclear-soluble and chromatin-bound protein fractions in response to DNA damage^16,17^. Using similar conditions, we isolated the nuclear-soluble and chromatin-bound fractions in response to increasing concentrations of PARG inhibitor (PARGi) and the DNA single-strand break inducing agent, methyl methanesulfonate (MMS) in the U2OS cell line. Upon treatment with the PARGi, PDD 00017273, we observed smearing above the PARP1 band, consistent with the increased accumulation of PAR chains on PARP1. In the presence of MMS and PARGi, we observed an increased proportion of PARP1 in the nuclear soluble fraction relative to the chromatin-bound fraction. [Figure 1A] To identify the subnuclear location of the increased PARP1, we performed laser microirradiation with PARP1 immunofluorescence. [Figure 1B] In response to laser-induced DNA damage, PARP1 is immediately recruited to the damage site and the signal resolves within 10 minutes after damage induction in the untreated condition. [Figure 1C, E] However, when cells were pre-treated with PARGi, we observed a distinct localization of PARP1 as foci throughout the nucleus 1 minute after laser-induced damage. We quantified the PARP1 intensity at the DNA damage stripe (as indicated by γH2AX signal) relative to total nuclear PARP1 and found that there was a significant decrease in the PARGi condition relative to the untreated condition. [Figure 1D] Consistent with previous reports, pre-treatment with the PARP inhibitor (PARPi) olaparib, led to a retention of PARP1 signal at the damage site relative to the untreated condition^18-20^. [Figure 1E, F] Strikingly, with PARGi pre-treatment, PARP1 had entirely reorganized as foci 10 minutes after damage induction. [Figure 1E] To rule out any off-target effects of the inhibitor, we used an siRNA to knockdown PARG and observed the same phenotype. [Supplementary Figure 1A, B, C] These results demonstrate that PARG inhibition reduces the chromatin binding of PARP1 owing to the accumulation of PARP1 as foci throughout the nucleus.

**Figure 1:**
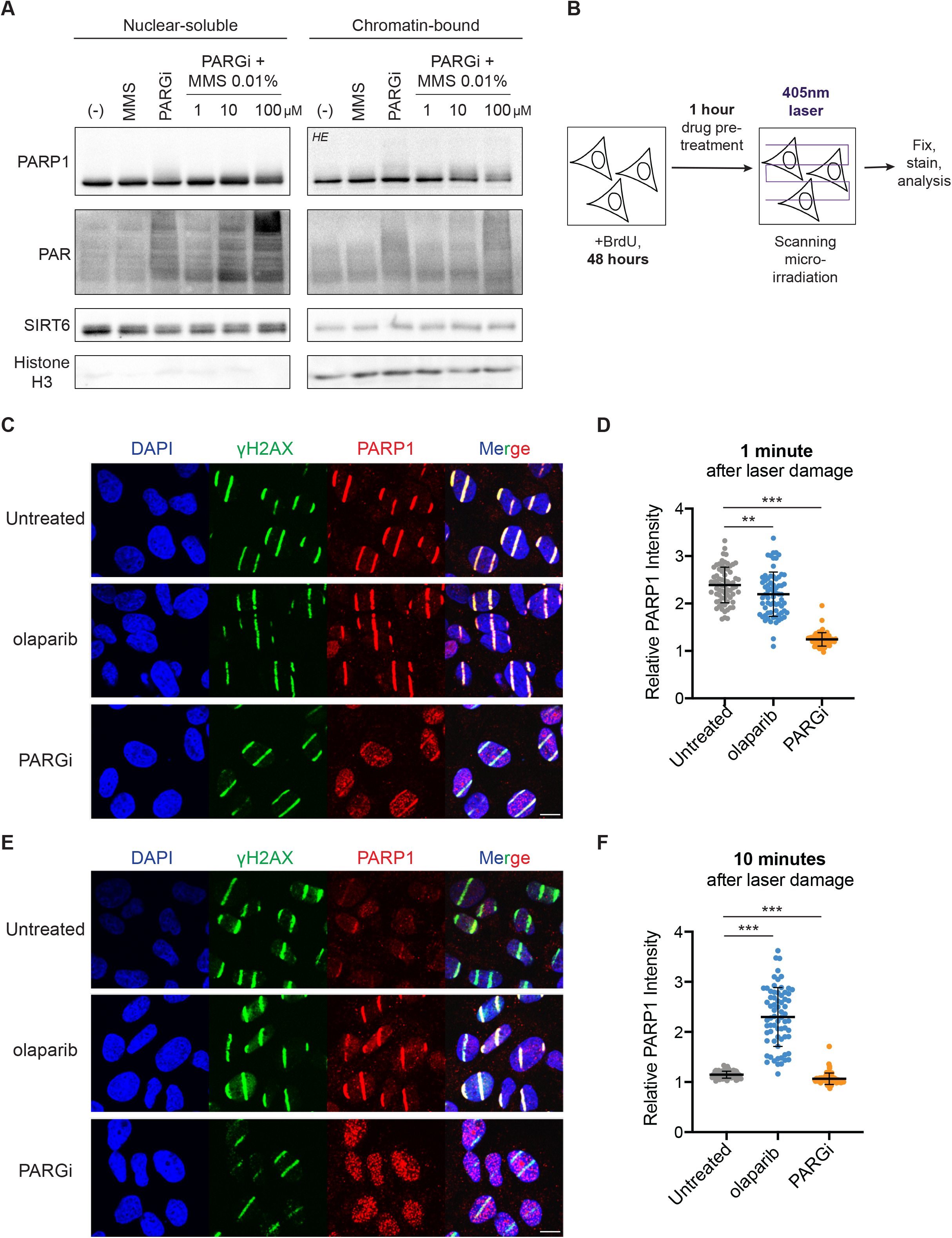
PARG inhibition reduces PARP1 chromatin binding through the accumulation of nuclear PARP1 foci. A) U2OS cells were treated as indicated and then subjected to subcellular fractionation. SIRT6 and Histone H3 represent the nuclear and chromatin loading controls respectively. An accumulation of PARylated PARP1 is visible in the last lane of the nuclear-soluble fraction with a concomitant decrease in the chromatin-bound fraction. HE = high exposure. B) Schematic overview of laser microirradiation experiments. C) Laser microirradiation was performed after pre-treatment with 5μM of the indicated drugs. Cells were fixed and stained for PARP1 and γH2AX at multiple time-points after laser-induced damage. Representative images at 1 minute (C) and 10 minutes (E) show PARP1 accumulating as foci in the PARGi condition. Scale bar = 10μm. D), F) Relative PARP1 intensity was calculated as the ratio of PARP1 signal at the γH2AX stripe normalized to background nuclear PARP1 signal in for all three conditions. **p< 0.01, ***p< 0.001 (two-tailed unpaired t-test), data presented as individual points with mean ± SD indicated, n ≥60 nuclei per condition).

### PARG inhibition leads to the formation of PARP1 aggregates

To monitor the formation of PARP1 foci with greater resolution, we developed a PARP1 live-cell imaging system in the U2OS cell line. We over-expressed PARP1 with a C-terminus enhanced GFP tag (PARP1-eGFP) in a PARP1-deficient background and confirmed PARP1 expression. [Figure 2A] We also confirmed that the over-expression of PARP1 re-sensitized U2OS to the PARPi, talazoparib. [Figure 2B] Using the PARP1-eGFP cell line, we performed similar laser microirradiation experiments in live cells. [Figure 2C] Briefly, using the near-UV 405nm laser, we made a 1μm^2^ bleach spot in PARP1-eGFP-expressing nuclei and monitored PARP1 recruitment and dissociation kinetics from the damage site. In untreated and PARGi-treated cells, PARP1 signal was visible as soon as 30 seconds after damage induction. [Figure 2D, E] In the untreated condition, PARP1 signal began to resolve 1 minute after damage induction. However, in the PARGi-treated condition, the PARP1 signal remained stable at the initial damage site up to 15 minutes. Additionally, similar to the fixed cell imaging findings, we observed that PARP1 localized as nuclear foci in response to PARGi pre-treatment. [Figure 2E] Using a custom imaging pipeline, we quantified the number of PARP1 foci over time. [Figure 2F] We observed that the PARP1 foci followed common protein aggregation kinetics starting with an initial nucleation phase (0-2 minutes), a rapid exponential growth phase (2-10 minutes) and a final plateau phase (10-15 minutes)^21^. These foci also maintained a constant size and intensity once formed, and persisted throughout the imaging leading us to classify them as PARP1 aggregates. [Figure 2G, Supplementary Movie 1]

**Figure 2:**
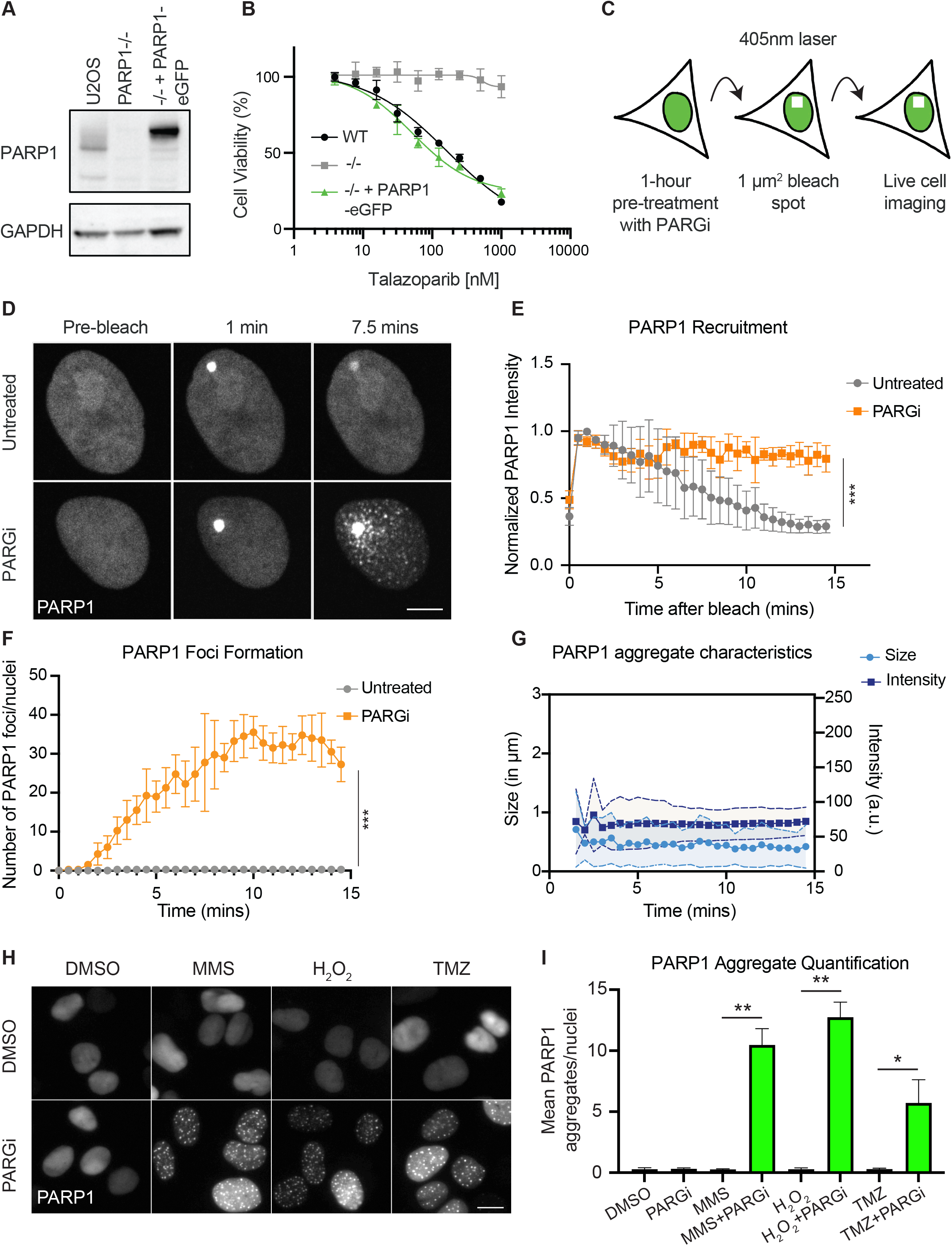
PARG inhibition leads to the formation of PARP1 aggregates in response to DNA damage. A) Western blot showing the overexpression of C-terminal GFP-tagged PARP1 in a U2OS PARP1-/-cell line. B) 5-day cell viability assay showing PARP1-eGFP overexpression re-sensitizes cells to talazoparib treatment. C) Schematic overview of live cell imaging to monitor PARP1 recruitment and dissociation in the PARP1-eGFP cell line. D) Cells were pre-treated with 10μM PARGi for 1 hour and subjected to laser-induced damage. Images were taken every 30 seconds for 30 frames and representative images are shown at pre-bleach, and 1- and 7.5-minutes post-bleach. E) Bright spot intensity was quantified and normalized to maximum PARP1 intensity across 4 different experiments and data are presented as mean ± SD. ***p< 0.001 (Welch’s two-sample t-test) F) PARP1 aggregates were quantified over time in the PARGi-treated and untreated condition. G) PARP1 aggregate size (in microns) was plotted on the left y-axis and aggregate intensity (in arbitrary units a.u.) was plotted on the right y-axis over time. Once PARP1 aggregates are formed and detected, their size and intensity remains stable. H) Representative images of PARP1-eGFP cells treated with the indicated treatments. Cells were treated with 0.01% MMS, 50μM H_2_O_2_ for 1 hour and 200μM TMZ for 2 hours in the presence and absence of 10μM PARGi. I) Mean PARP1 aggregates were quantified across 2 independent experiments. *p< 0.05, **p< 0.01 (two-tailed unpaired t-test), data presented as individual points with mean ± SD indicated, n ≥70 nuclei per condition. Scale bar = 10μm.

To rule out the possibility that this phenotype was specific to laser-induced damage, we treated the U2OS PARP1-eGFP cells with DNA damaging agents including MMS, hydrogen peroxide (H_2_O_2_), and temozolomide (TMZ). All three agents produce a proportion of oxidative and/or alkylating DNA damage, leading to the creation of abasic sites, which trigger PARP1-mediated DNA repair^22^. The cells were concomitantly treated with PARGi and showed a similar PARP1 aggregation as seen in the laser-induced fixed and live cell imaging. [Figure 2H] In comparison, no PARP1 aggregation was detected in the untreated or single agent conditions. [Figure 2H, I] Similar to the laser microirradiation experiments, we observed that the PARP1 aggregates accumulated at sites distinct from the site of damage as indicated by the 0% co-localization between PARP1-eGFP and γH2AX and 53BP1 staining. [Supplementary Figure 2A, B] These findings demonstrate that PARGi treatment with DNA damage causes nuclear PARP1 aggregation.

### PARP1 aggregates contain PARylated PARP1

Given that the PARP1 aggregates only form in conditions treated with PARGi, we hypothesized that these aggregates contained PARylated PARP1. To enable to the detection of PAR in live cells, we created a PARP1/PAR dual sensor cell line. Briefly, we adapted a method utilizing the PAR binding domain of APLF to visualize PAR chains and cloned it into an mCherry-expressing plasmid^23,24^. We co-nucleofected the APLF-mCherry and PARP1-eGFP plasmids into a PARP1-/-U2OS cell line and sorted out double positive cells. To validate the specificity of the sensor, we pre-treated cells with the PARPi, olaparib, to abrogate PAR chain synthesis. Consistent with this, we observed a complete disappearance of PAR chain signal at the damage site despite PARP1 recruitment. [Figure 3A, B] Using this cell line, in the presence of PARGi pre-treatment, we observed PARP1 aggregation as expected. [Figure 3C] Interestingly, the PAR signal continued to increase over time and entirely co-localized with the signal from the PARP1 aggregates. [Figure 3C] This revealed that the PARP1 aggregates contained PAR chains. To test whether the PAR chains on PARP1 were essential for the formation of these aggregates, we identified two catalytic variants of PARP1 with defective autoPARylation. Mutating the glutamate at the 988 position in PARP1 to alanine and aspartate has previously been reported to either completely abrogate or significantly diminish PAR chain synthesis respectively by PARP1 *in vitro*^25^. We cloned these PARP1 catalytic variants into the eGFP plasmid, selected and sorted GFP+ cells, and validated their PARP1 and PAR expression. [Figure 3D] As expected, the two PARP1-E988D clones showed reduced PAR chain synthesis at baseline. The PARP1-E988A clone showed no PAR chain synthesis, however, the single cell clones with this mutation were difficult to maintain. This is possibly due to the extreme toxicity of this mutation to cells as previously described^26^. Using both PARP1 catalytic variant cell lines, we performed laser microirradiation and live cell imaging and found a complete abrogation of PARP1 aggregate formation in response to PARGi treatment. [Figure 3E, G] Interestingly, inhibiting the catalytic activity of PARP1 also abrogated PARP1 accumulation at the original damage site in response to PARGi treatment. [Figure 3F] These results demonstrate that PAR chains are present in these PARP1 aggregates and that PARP1 catalytic activity is necessary for the formation of these aggregates.

**Figure 3:**
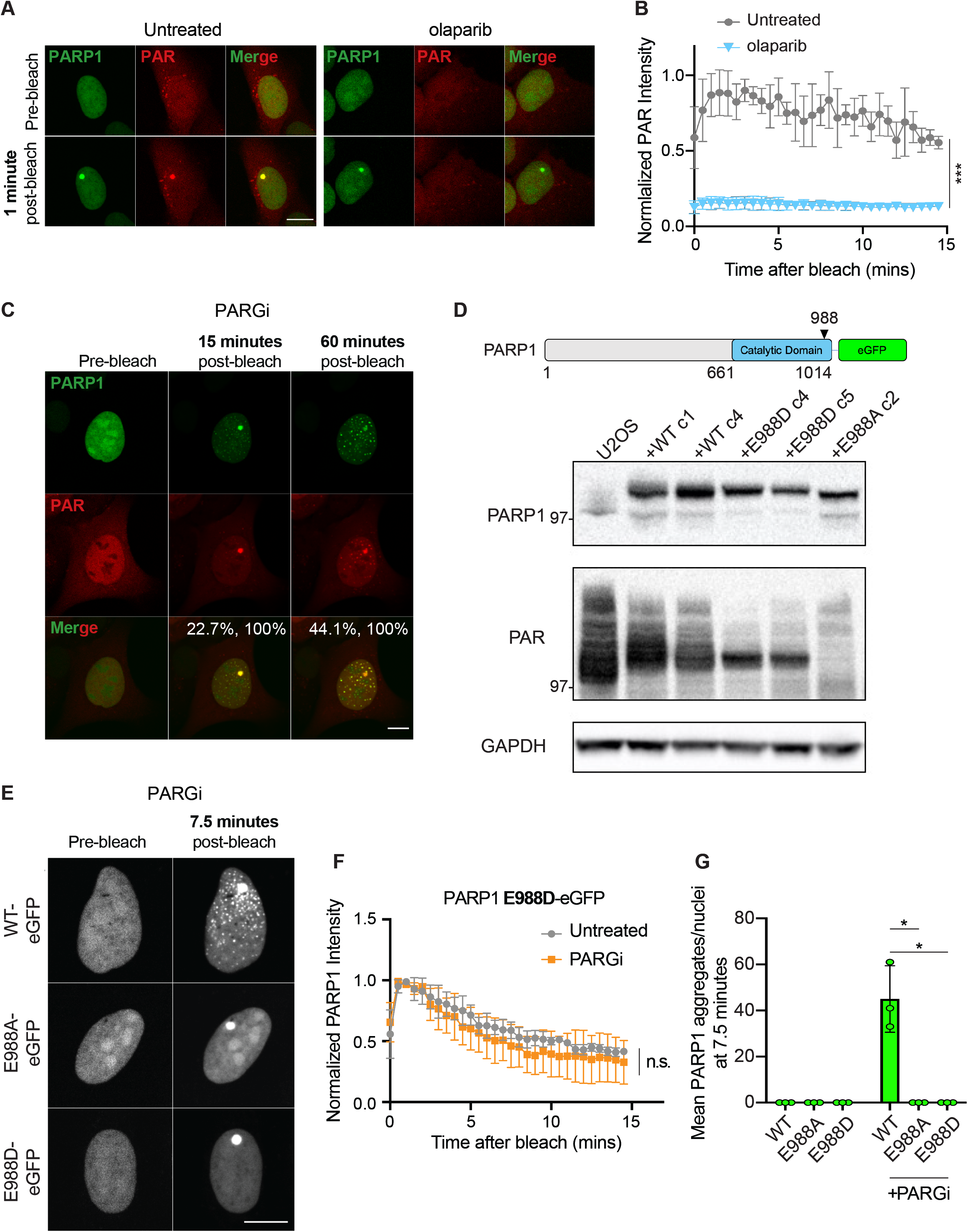
PARP1 catalytic activity is required for the formation of PARP1 aggregates in response to DNA damage. A) Representative images of the validation of the PAR sensor cell line with olaparib treatment. Upon olaparib treatment, no PAR sensor signal is detected 1 minute after laser-induced damage. B) Bright spot intensity was quantified and normalized to maximum PAR intensity across 4 different experiments, ***p< 0.001 (Welch’s two-sample t-test). (C) Representative images of PARP1-eGFP, APLF-PBZ-mCherry, and the merged images taken pre-bleach, and 15-minutes and 60 minutes after laser-induced damage. Cells were pre-treated with 10μM PARGi for 1 hour. The colocalization of PARP1 with PAR and PAR with PARP1 signal were represented as percentages calculated using ImageJ with a tolerance of 1 pixel. D) Schematic of full-length PARP1 protein showing the catalytic domain and the site of the point mutation. Western blot showing overexpression of PARP1 catalytic variants, E988D, producing medium PAR chains, and E988A, producing no PAR chains in the PARP1-/-cell line. The PARP1 catalytic variants show reduced PAR chain expression compared to PARP1 wildtype (WT). E) Representative images of PARP1-WT, PARP1-E988A, E988D-eGFP cells pre-bleach and 7.5 minutes post-bleach after pre-treatment with 10μM PARGi for 1 hour. F) Quantification of normalized PARP1 intensity at the site of damage with and without PARGi treatment after laser-induced damage over 15 minutes. G) Quantification of PARP1 aggregates in PARP1-WT, E988A, E988D-eGFP cell lines with and without PARGi treatment at 7.5 minutes after laser-induced damage. PARP1 aggregates form only in the PARP1 catalytically-proficient cell line. Data are presented as mean ± SD, *p< 0.05 (two-tailed unpaired t-test), scale bar = 10μm.

### PARP1 aggregates are associated with increased cell death

Given that the location of these PARP1 aggregates remained relatively stable over time, we next investigated whether these PARP1 aggregates persisted long-term. To test this, we identified a low dose of MMS, which could be used to perform imaging over 24 hours without significantly affecting cell viability. We then treated cells with low dose MMS in combination with PARGi and confirmed that PARP1 aggregates also formed and persisted under these conditions at 24 hours. [Figure 4A] Interestingly, we observed in the 80% of cells that contained nuclear PARP1 aggregates, approximately 70% of those cells also contained cytoplasmic PARP1 signal. [Figure 4B] It is well established that cleaved PARP1 is translocated to the cytoplasm to activate cell death signaling cascades^27^. Since 100% of cytoplasmic PARP1 signal was connected to nuclear PARP1 aggregation, we hypothesized that the aggregates could be promoting cytotoxicity. We performed a cell cycle analysis on cells treated with the same conditions for 24 hours. [Figure 4C] In response to MMS-induced damage, cells are arrested in the G2 phase to allow for the repair of the DNA damage prior to mitosis^28^. However, in the PARGi and MMS-treated condition, we observed a loss of the expected G2 stalling and a proportional increase in the sub-G1 population, indicating greater DNA fragmentation associated with cell death in this condition. [Figure 4D] DNA fragmentation and cytoplasmic PARP1 are both hallmarks of apoptosis, however, when the cells were treated with PARGi and MMS, along with a pan-caspase inhibitor, Z-VAD-FMK, we did not observe a rescue of cytotoxicity, likely pointing to a non-apoptotic form of cell death. [Figure 4E] We then tested whether the combination of both MMS and PARGi would lead to increased toxicity compared to either single agent over a 5-day cell viability assay. MMS showed synergistic cell kill with PARGi in the U2OS PARP1-eGFP cell line. [Figure 4F] However, the PARP1 catalytic-deficient cell lines did not show a similar synergistic cell killing confirming that the PARP1 aggregation phenotype is associated with increased toxicity of PARGis. [Figure 4G, H] This demonstrated that the persistent PARP1 aggregates were associated with increased non-apoptotic cell death signaling and contributing to the toxicity associated with PARGis.

**Figure 4:**
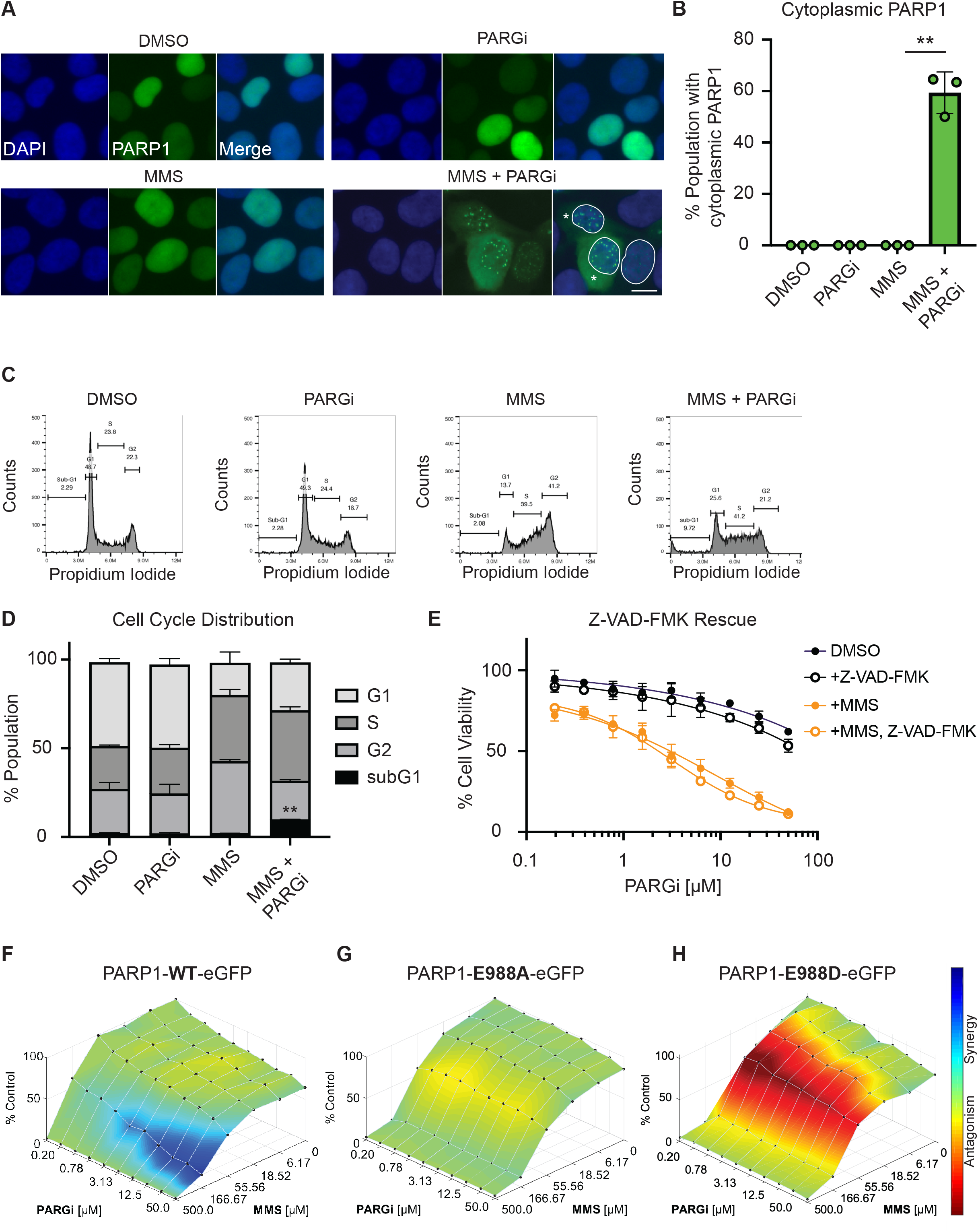
PARP1 aggregates are associated with increased cell death. A) Representative images of PARP1-eGFP in response to the indicated damaging treatments. Cells were treated with 0.01% MMS and 10μM PARGi for 24 hours. Diffuse cytoplasmic PARP1 signal (indicated with an asterisk) is observed only in the MMS + PARGi condition and in cells with nuclear PARP1 aggregates. B) Percentage of cells containing cytoplasmic PARP1 was quantified across the treatment conditions. Data are presented as mean ± SD, **p< 0.01 (two-tailed unpaired t-test) C) Cell cycle distribution analysis was performed on U2OS PARP1-eGFP cells DMSO, 0.01% MMS, 10μM PARGi, and 0.01% MMS and 10μM PARGi for 24 hours. Representative histograms of all four conditions are shown. D) Quantification of the different cell cycle stages shows an accumulation of a sub-G1 population in the combination treatment. Data are presented as mean ± SD, **p< 0.01 (two-tailed unpaired t-test) E) Cells were treated with 0.0125% MMS and increasing concentrations of PARGi with and without 50μM Z-VAD-FMK (pan-caspase inhibitor) for 4 days. Percent cell viability was plotted as mean ± SD. E) U2OS PARP1-eGFP cell line was treated with increasing concentrations of PARGi and MMS for 5 days. Results were plotted using the Loewe synergy model in the Combenefit software. A synergistic cell kill was seen with the combination. F) Similar cell viability assays were performed in the E988A-eGFP and E988D-eGFP (G) cell lines. No synergy was visible in these cell lines under the same treatment conditions. Scale bar = 10μm.

Finally, we tested the PARP1 aggregation phenotype in a clinically relevant cell line, OVCAR-8, a high-grade ovarian cancer cell line with a BRCA1 hypermethylation. We generated an OVCAR-8 PARP1-eGFP cell line and validated PARP1 protein expression and re-sensitization to talazoparib treatment. [Supplementary Figure 3A, B] Similar to the U2OS cell line, we observed PARP1 aggregate formation in response to PARGi and DNA damaging agents inducing PARP1 activation. [Supplementary Figure 3C, D] Additionally, these PARP1 aggregates also formed at sites distinct from sites of damage marked by γH2AX. [Supplementary Figure 3E] PARGi itself was not able to cause substantial cell death in these cell lines, however, the combination of PARGi and MMS led to synergistic cell kill in the OVCAR-8 cell line. [Supplementary Figure 3F]

## Discussion

Here, we report a novel mechanism of action of PARG inhibitors through the formation of cytotoxic, nuclear PARP1 aggregates. PARG inhibitors have been shown to cause increased PARylation of PARP1 however, how this affects DNA repair, replication, etc. has yet to be elucidated. Our results corroborate previous findings in *in vitro* and cell-based systems that an increased accumulation of PAR chains on PARP1 (in this study through PARG inhibition), reduces the chromatin binding of PARP1^4,11,29^. This represents the first report to observe that this accumulation of PAR chains on PARP1 results in the widespread nuclear aggregation of PARP1. Previous reports likely failed to visualize this aggregation owing to technical limitations associated with PARP1 immunofluorescence and the use of laser microirradiation to monitor short-term protein chromatin-recruitment rather than long-term protein chromatin-dissociation kinetics. Our data demonstrates that these aggregates are formed through interactions between PARylated PARP1. Previous literature has demonstrated that PAR chains represent ideal scaffolds for the formation of large-scale protein complexes or biomolecular condensates owing to their high charge density and low complexity^30,31^. Specifically, PAR chains have been shown to accelerate the formation of protein aggregates of intrinsically disordered proteins *in vitro*^32,33^. This property of PAR chains has been shown to organize DNA repair hubs, stress granules, and processes within nuclei, to name a few^31^. Given our findings, we speculate that in these instances PARylated PARP1 could be organizing these transient biomolecular structures.

The most striking feature of these PARP1 aggregates is their formation at sites distinct from the site of damage. Our results demonstrate that these aggregates do not co-localize with proteins recruited to damage sites such as 53BP1, and are likely not bound to chromatin. One hypothesis for where these aggregates could be accumulating is based on multiple reports of PARP1 studies in *Drosophila*. These reports had shown that following auto-modification, PARylated PARP1, bound to other nuclear proteins, migrated from the chromatin to Cajal bodies in the nucleus^34^. While the specific function of this translocation is still under study, one hypothesis is that PARylated PARP1 mediates the recruitment of RNA-related proteins that need to be localized to Cajal bodies. The other hypothesis is that PARylated PARP1 could be transported into Cajal bodies to allow for the efficient recycling of PARP1 and PAR chains following repair. PAR chains have previously been shown to be substrates for the generation of nuclear ATP in conditions of high ATP requirement such as chromatin relaxation in response to DNA repair^35^. HyperPARylated PARP1 could normally be sequestered away into Cajal bodies to allow for the recycling of ADP-ribose and PARP1. However, in the case of PARG inhibition, this recycling pathway would be severely overwhelmed causing the persistence of these PARP1 aggregates over 24 hours.

PARG knockdown, similar to PARP1 knockdown, has previously been shown to delay the rate single strand break repair^12,36^. Our results support this observation as the accumulation of PARP1 as aggregates would prevent its recycling to respond to other DNA lesions, thereby delaying resolution of damage sites. Another study has shown that PARP1 scans the genome in search for DNA lesions through a monkey-bar mechanism^37^. Excessive PARylation on PARP1 would also impede this scanning and genome maintenance function of PARP1. Additionally, with persistent PAR chains, any PAR-mediated repair complexes formed to respond to DNA damage sites would be delayed in their disassembly and reassembly at other damage sites. Since these aggregates form at sites distant from the original site of damage, this could lead to the mis-localization of proteins recruited in response to PAR such as X-ray repair cross-complementing protein 1 (XRCC1), Ligase 3, etc. Additionally, a number of DNA repair proteins have been shown to get recruited to DNA damage in response to PAR chain chromatin relaxation as opposed to the binding of these proteins to PAR chains^38^. Increased retention of PAR chains at the chromatin in response to PARGi treatment may also drive the accumulation of DNA-binding proteins to DNA with no functional role in DNA repair leading to overcrowding at these DNA lesions. In all these scenarios, PARG inhibition slows down DNA repair and increases the proportion of unresolved DNA damage.

Finally, our cell viability assays conclude that conditions where PARP1 aggregates are formed are associated with greater levels of cell toxicity than PARGi or DNA damage alone. While the precise molecular mechanisms that cause PARP1 aggregation to trigger the cell death cascade are still unclear, cleaved cytoplasmic PARP1 is considered to be a hallmark of cell death. Our results demonstrate a cytoplasmic PARP1 signal and increased DNA fragmentation is associated with nuclear PARP1 aggregation. However, an apoptosis inhibitor, Z-VAD-FMK, was unable to rescue the cell death association with PARGi and DNA damage treatment. This could point to Future work will identify the whether the aggregates directly or indirectly activate a non-apoptotic form of cell death. Overall, this report sheds light on a novel mechanism of PARGi-mediated toxicity through the formation of persistent, nuclear PARP1-aggregates.

## Materials and Methods

### Cell lines and culture conditions

U2OS cells were obtained from ATCC and cultured in high glucose Dulbecco’s modified Eagle’s medium (Thermo Fisher Scientific) with L-glutamine containing 10% fetal bovine serum (Sigma-Aldrich) and 1% Penicillin Streptomycin (Thermo Fisher Scientific). OVCAR-8 cells were obtained from the DCTD Tumor Repository at the National Cancer Institute. The cells were cultured in Roswell Park Memorial Institute (RPMI)-1640 with L-glutamine (Thermo Fisher Scientific) with 10% fetal bovine serum and 1% Penicillin Streptomycin. All cells were maintained at 37°C with 5% CO_2_.

### Chemicals and reagents

The following were purchased from commercial sources as indicated: dimethyl sulfoxide (Sigma-Aldrich), methyl methanesulfonate (Sigma-Aldrich), PDD 00017273 (PARGi, Tocris), olaparib (MedChem Express), 5-Bromo-2’-Deoxyuridine (Sigma Aldrich), talazoparib (Selleckchem), hydrogen peroxide (Sigma-Aldrich), and temozolomide (Selleckchem).

### Western blot

Pellets were subjected to subcellular fractionation according to manufacturer’s instructions (Thermo Scientific). Protein lysates were quantified with Bradford reagent (Bio-Rad) and depending upon the protein of interest, 25-40 μg of lysate was loaded on to NuPAGE 4 to 12%, Bis-Tris gels (Invitrogen). Proteins were transferred on to Immun-Blot PVDF membranes (Bio-Rad) using the Mini Trans-Blot Cell (Bio-Rad). Membranes were blocked in 5% BSA in TBST for 1.5 hours and primary antibodies were added overnight at the following concentrations: anti-PARP1 (Cell Signaling Technology, 1:1000), anti-PAR (Millipore, 1:5000), anti-SIRT6 (Cell Signaling Technology, 1:1000), anti-Histone H3 (GeneTex, 1:1000), anti-GAPDH (Proteintech, 1:1000), anti-PARG (Millipore, 1:500). Secondary antibodies were added at a dilution of 1:5000 in TBST and incubated for 1.5 hours. Protein levels were visualized using Clarity ECL substrate (Bio-Rad) and images were acquired on the Chemidoc Imaging System (Bio-Rad).

### Laser microirradiation and immunofluorescence

Laser microirradiation experiments were performed as previously described with the Leica SP8 scanning confocal microscope equipped with a 405nm diode^39^. Immediately after microirradiation, cells were fixed and permeabilized in 3% PFA, 8% sucrose, 0.5% Triton X in PBS for 15 minutes, washed, and blocked in 5% goat serum, 5% fetal bovine serum, 0.2% Triton-X in PBS overnight at 4C. The following day primary antibodies were added to the slide and the day after, secondary antibodies were added at a 1:500 dilution and DAPI was added at 1:100. Coverslips were mounted on glass slides using fluorescence mounting medium (Dako). Slides were imaged using the TiE inverted spinning disc confocal microscope (Nikon). Images were analyzed using the “Stripenator” pipeline. The relative PARP1 intensity was calculated as the intensity of PARP1 at the stripe area versus intensity at the non-stripe area marked by γH2AX signal. Heat maps were generated for certain repair proteins using the average protein intensity at the stripe relative to the background nuclear signal.

### siRNA transfection

U2OS cells were reverse transfected with PARG siRNA (Horizon Discovery, L-011488-00-0005) and a non-targeting control. siRNAs were resuspended in 1X siRNA buffer (Horizon Discovery). Using 1μL of the diluted siRNA, 250,000 cells were reverse transfected using RNAimax (Thermo Fisher Scientific) in OptiMEM (Thermo Fisher Scientific). Protein knockdown was assessed using Western Blot 72 hours after transfection.

### Generation of PARP1-eGFP cell lines

PARP1 knockout cell lines were generated in the U2OS and OVCAR8 cell line using lentivirus as described before^40^. Knockouts were validated by western blotting for PARP1 expression and Sanger sequencing. The C-terminal-tagged PARP1-eGFP was generously provided by Dr. Shan Zha at Columbia University. A sgRNA-resistant PARP1 cDNA sequence was generously provided by Dr. Ryan Jensen at Yale University and cloned into the PARP1-eGFP plasmid. All sequences were validated using Plasmidsaurus sequencing. 2μgs of the PARP1-sgRNAR-eGFP plasmid was nucleofected into the U2OS PARP1-/-and OVCAR8 PARP1-/-cell line using the Nucleofector (Lonza). Cells were selected in 2μg/mL G418 for 10 days and GFP-positive cells were sorted and used for subsequent experiments. The PARP1 variant cDNA sequences were obtained as gene blocks (Genscript) and were cloned and nucleofected using a similar strategy. The PAR sensor cell line was generated by cloning the APLF cDNA sequence (Genscript) into the pHR-FUSN-mCh-CRY2WT plasmid (Addgene; 101223). The PARP1-eGFP plasmid and APLF-mCherry plasmid were co-nucleofected into the U2OS PARP1-/-cell line and selected with 2μg/mL G418 for 10 days. mCherry- and GFP-positive cells were sorted and used for subsequent experiments.

### Cell viability assays

Cells were seeded at 1000 cells/well in a 96-well plate and were treated 24 hours later with the indicated drug concentrations. 5 days later, cells were fixed in 4% formaldehyde, washed with PBS, and stained with 1μg/mL Hoechst 33342 (Thermo Fisher Scientific). Plates were imaged using the Cytation 3 (BioTek) and cells were counted using CellProfiler. For synergy assays, cells were plated and analyzed similarly, and synergy was calculated using the Loewe method in Combenefit^41^.

### Live cell imaging and data analysis

All live cell imaging was performed in either the PARP1-eGFP cell lines or the PAR sensor cell line. Cells were seeded in 8-well imaging chambers (Miltenyl Biotec) and their media was changed to phenol-red free imaging media (Thermo Fisher Scientific) prior to imaging. Cells were pre-treated for 1 hour with either 10μM PARGi or 5μM olaparib. Laser microirradiation was performed by bleaching a 1 μm^2^ bleach spot with the 405nm diode laser at 100% laser intensity, speed 100Hz, for 1 iteration. Images were taken every 30 seconds for 30 frames.

For PARP1 aggregate analysis, cell images were first processed to detect bright spots in the nucleus, and then these spots were tracked as trajectories across multiple frames of the image. Using the scikit-image package in Python, the spot detection algorithm applies a white-tophat transform to the frame before applying the Canny edgefinding algorithm^42,43^. Each resulting edge within the nucleus is identified as a bright spot if the edge is a closed polygon. Tracking of each detected bright spot was performed using the link() function of the trackpy package in Python, with parameter search_range=3 and memory=3^44^. Trajectories were further filtered to only those containing spots for at least 4 frames using filter_stubs() with parameter threshold=4. Spot location, spot size, and mean spot intensity were computed and tracked for each resulting trajectory. For PARP1 and PAR signal in the PAR sensor cell line, co-localization analysis was performed using the Focinator pipeline. A tolerance of 1 pixel was set for co-localization analysis. Percent co-localization was calculated for every nucleus and then averaged across independent biological experiments.

### Immunofluorescence

Cells were seeded in 8-chamber slides (Millipore) and treated with the indicated drug concentrations. Cells were fixed with 4% paraformaldehyde (Electron Microscopy Sciences) in PBS for 10 minutes, washed, and then permeabilized with 0.05% Triton-X for 10 minutes. Slides were blocked in 10% goat serum in 0.02% Triton-X in PBS and antibodies were added overnight at 4°C at the following concentrations: anti-PARP1 (R&D Biosystems, 1:1000), anti-γH2AX (Millipore, 1:1000), anti-53BP1 (Novus Biologicals, 1:1000). For PARP1-eGFP immunofluorescence, cells were fixed, permeabilized, and immediately mounted using Vectashield antifade mounting medium with DAPI (Vector Laboratories). Slides were imaged using the Keyence BZ-X800 using the 40X objective. PARP1-eGFP, γH2AX, and 53BP1 foci were quantified using the Focinator pipeline.

### Flow cytometry

Cells were seeded in 60mm dishes and treated with the indicated drug concentrations. After 24 hours, cells were harvested and fixed drop-wise in 70% ice-cold ethanol for 15 minutes. After washing with PBS, cells were stained with Propidium Iodide/RNase A (BD Biosciences) for 15 minutes in the dark. Stained cells were then run on the CytoFlexLX and the data was analyzed using FlowJo software.

### Quantification and statistical analysis

All statistical analysis was performed using the two-tailed Student’s *t*-test or Welch’s sample t-test in Prism (GraphPad). In all cases n.s. indicates not significant, *p< 0.05, **p< 0.01, ***p< 0.001.

## Acknowledgements

We thank the Yale West Campus Imaging Core and for the support and assistance in this work. We also thank Yale Flow Cytometry for their assistance with the cell sorting service.

