## Supplementary Figures 1-3 for "PARG inhibition induces nuclear aggregation of PARylated PARP1"

**Supplementary Figure 1**: PARG knockdown produces PARP1 foci in response to laser-induced DNA damage.

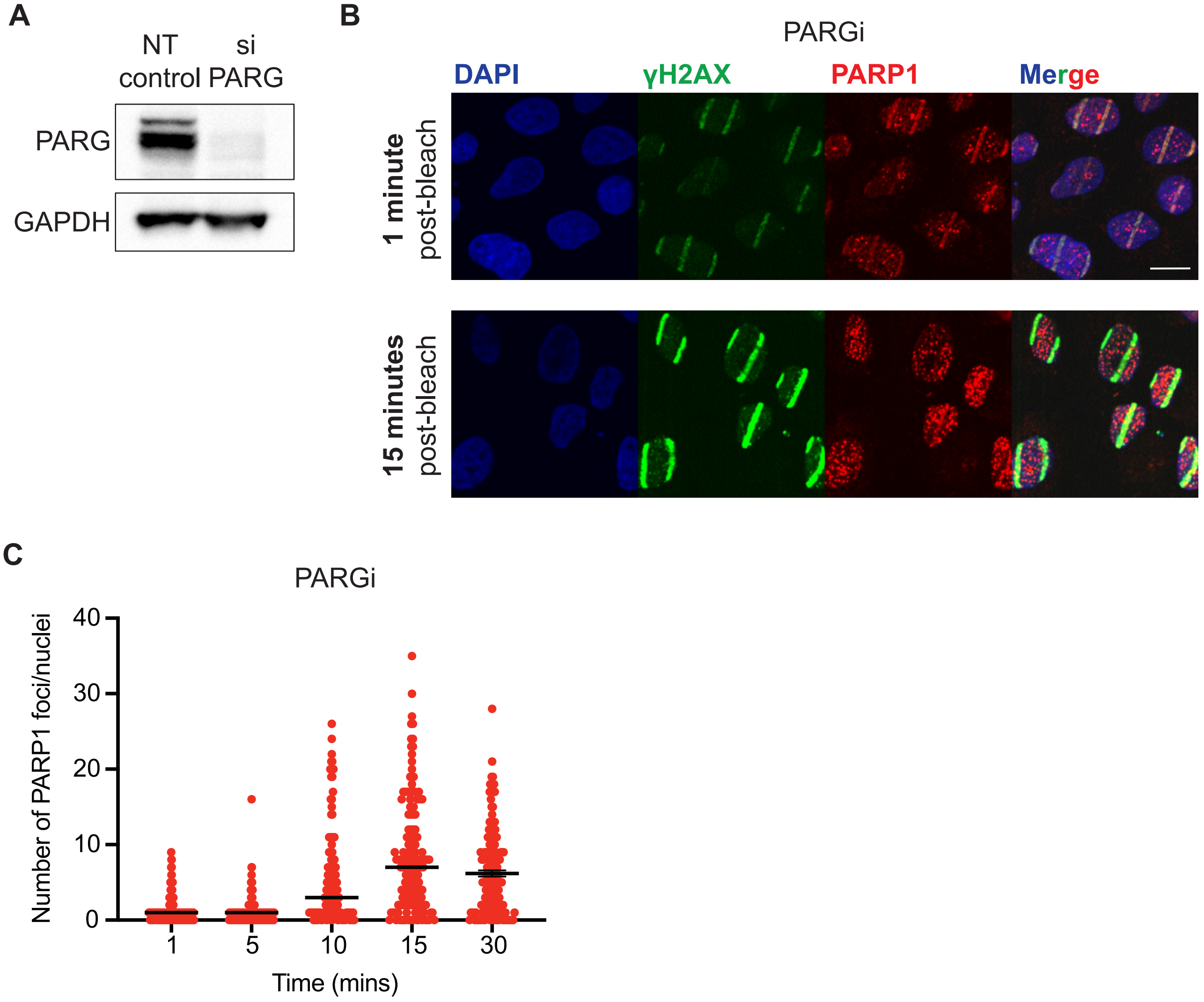

**Supplementary Figure 1: PARG knockdown produces PARP1 foci in response to DNA damage.** A) Western blot showing reduced PARG expression levels after siRNA knockdown of PARG relative to the non-targeting control. B) Laser microirradiation was performed followed by fixed cell imaging in the PARG knockdown and control condition. Representative images at 1 and 15-minutes post-bleach are presented. siPARG condition shows nuclear PARP1 foci at sites distant from the original site of damage as indicated by the γH2AX stripe. C) Quantification of number of PARP1 foci per nuclei in the siPARG condition over time. Mean is indicated and >70 nuclei were quantified for each timepoint.

**Supplementary Figure 2:** PARP1 aggregation occurs at sites distinct from the sites of damage

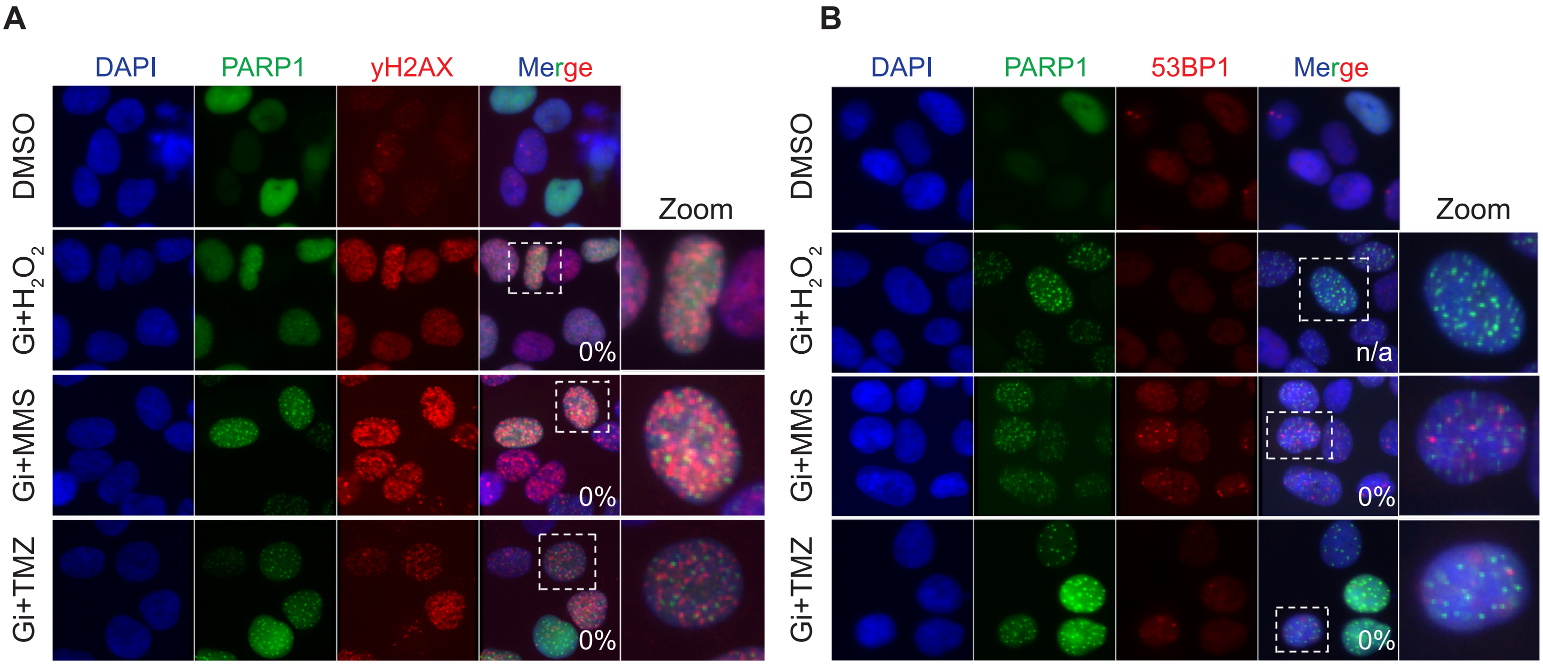

**Supplementary Figure 2: PARP1 aggregation occurs at sites distinct from the sites of damage.** A) Representative images of PARP1-eGFP cells treated with the indicated DNA damaging agents. Cells were treated with 0.01% MMS, 50μM $H_2O_2$ for 1 hour and 200μM TMZ for 2 hours in the presence and absence of 10μM PARGi. Slides were fixed and stained for γH2AX and 53BP1 (B). Co-localization analysis of PARP1-eGFP with γH2AX and 53BP1 was performed and indicated in the bottom right corner of the images. Scale bar = 10μm.

**Supplementary Figure 3**: PARP1 aggregates form in response to DNA damage and PARGi treatment in the OVCAR-8 cell line.

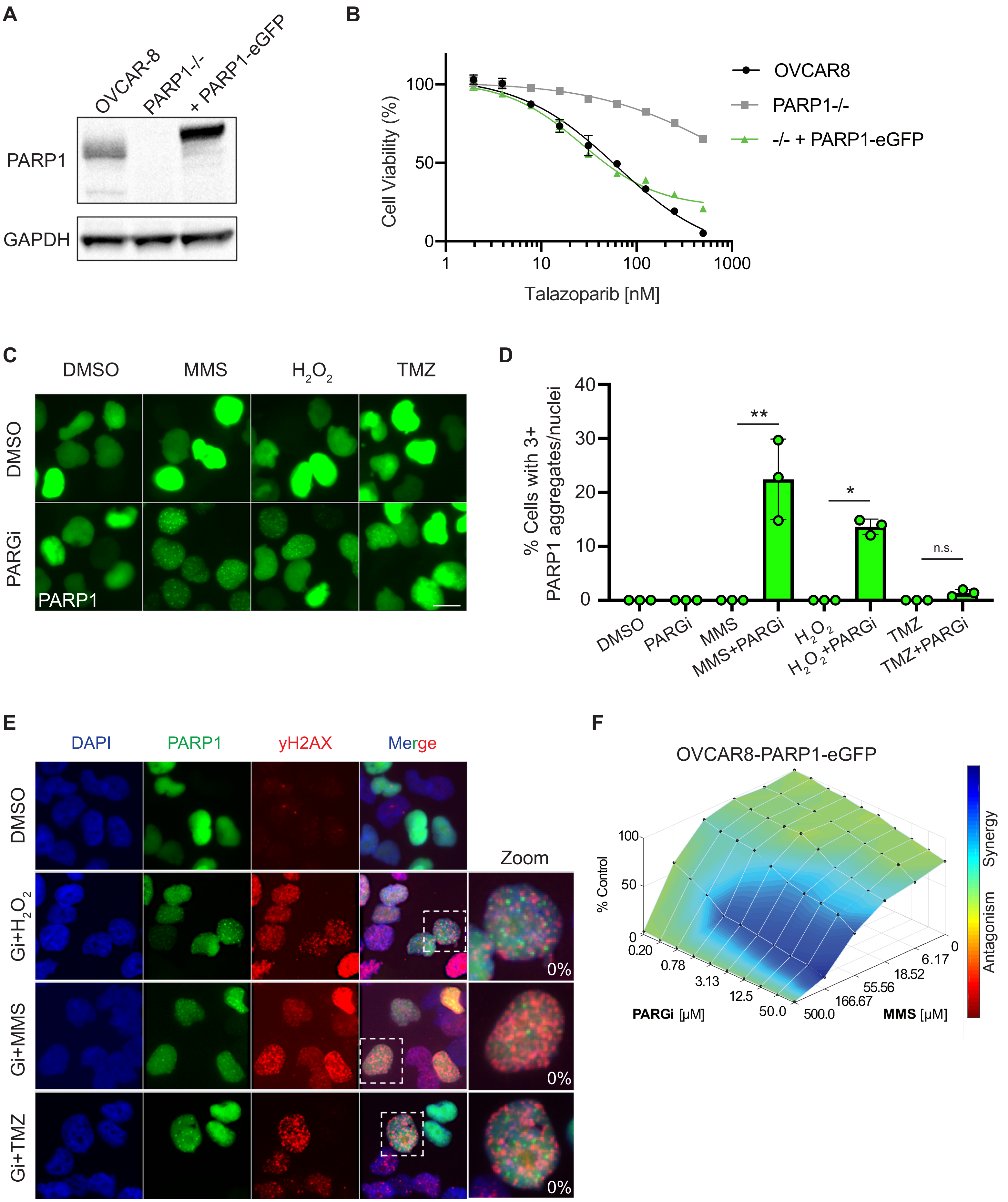

**Supplementary Figure 3: PARP1 aggregates form in response to DNA damage and PARGi treatment in the OVCAR-8 cell line.** A) Western blot showing the overexpression of C-terminal GFP-tagged PARP1 in an OVCAR-8 PARP1-/- cell line. B) 5-day cell viability assay showing PARP1-eGFP overexpression re-sensitizes cells to talazoparib treatment. C) Representative images of PARP1-eGFP cells treated with the indicated treatments. Cells were treated with 0.01% MMS, 50µM $H_2O_2$ for 1 hour and 200µM TMZ for 2 hours in the presence and absence of 10µM PARGi. D) Percent nuclei with greater than 3 PARP1 aggregates were quantified. *$p < 0.05$, **$p < 0.01$ (two-tailed unpaired t-test). Data presented as individual points with mean ± SD indicated, n ≥70 nuclei per condition. E) Representative images of OVCAR-8 PARP1-eGFP cells treated the same as in C. Slides were additionally stained for γH2AX. Co-localization analysis of PARP1-eGFP with γH2AX was performed and indicated in the bottom right corner of the images. F) OVCAR-8 PARP1-eGFP cell line was treated with increasing concentrations of PARGi and MMS for 5 days. Results were plotted using the Loewe synergy model in the Combenefit software. A synergistic cell kill was seen with the combination. Scale bar = 10µm.

**Supplementary Movie 1:** U2OS PARP1-eGFP cells were pre-treated with 10μM PARGi for 1 hour and bleached with the 405nm laser at time t=0. Images of the same nuclei were taken every 30 seconds for up to 15 minutes. The movie shows PARP1 aggregates forming and persisting throughout the nucleus in response to laser-induced damage.
